# Three stages in the development of the cyst wall of the eye pathogen *Acanthamoeba castellanii*

**DOI:** 10.1101/534487

**Authors:** Pamela Magistrado-Coxen, Yousuf Aqeel, A Lopez, John Samuelson

## Abstract

When deprived of nutrients, trophozoites of the eye pathogen *Acanthamoeba castellanii* make a cyst wall, which contains cellulose and has two layers connected by cone-shaped ostioles. We recently showed chitin is also present and identified three sets of lectins, which localize to the ectocyst layer (Jonah lectin) or the endocyst layer and ostioles (Luke and Leo lectins). To determine how the cyst wall is made, we examined encysting protists using structured illumination microscopy, probes for glycopolymers, and tags for lectins. In the first stage (3 to 9 hr), cellulose, chitin, and a Jonah lectin were each made in dozens of encystation-specific vesicles. In the second stage (12 to 18 hr), a primordial wall contained both glycopolymers and Jonah lectin, while small, flat ostioles were outlined by a Luke lectin. In the third stage (24 to 36 hr), an ectocyst layer enriched in Jonah lectin was connected to an endocyst layer enriched in Luke and Leo lectins by large, conical ostioles. Jonah and Luke lectins localized to the same places in mature cyst walls (72 hr) independent of the timing of expression. The Jonah lectin and the glycopolymer bound by the lectin were accessible in the ectocyst layer of mature walls. In contrast, Luke and Leo lectins and glycopolymers bound by the lectins were mostly inaccessible in the endocyst layer and ostioles. These results show that cyst wall formation is a tightly choreographed event, in which glycopolymers and lectins combine to form a mature wall with a protected endocyst layer.

**Importance:** While the cyst wall of *Acanthamoeba castellanii,* cause of eye infections, contains cellulose like plants and chitin like fungi, it is a temporary, protective structure, analogous to spore coats of bacteria. We showed here that, unlike plants and fungi, *A. castellanii* makes cellulose and chitin in encystation-specific vesicles. The outer and inner layers of cyst walls, which resemble the primary and secondary walls of plant cells, respectively, are connected by unique structures (ostioles) that synchronously develop from small, flat circles to large, conical structures. Cyst wall proteins, which are lectins that bind cellulose and chitin, localize to inner or outer layers independent of the timing of expression. Because of its abundance and accessibility in the outer layer, the Jonah lectin is an excellent target for diagnostic antibodies. A description of the sequence of events during cyst wall development is a starting point for mechanistic studies of its assembly.

## Introduction

Infections of the cornea (keratitis), which may lead to scarring and blindness, are caused by bacteria (e.g. *Neisseria trachomatis*), fungi (e.g. *Fusarium sp*.), or protists (*Acanthamoeba castellanii*) (the focus here) (1–3). In immunocompetent persons, *Acanthamoeba* infections are rare but difficult to treat (4, 5). In immunosuppressed patients, *Acanthamoebae* may cause encephalitis (6). *Acanthamoeba* is an emerging pathogen, because 80 to 90% of infections are associated with growing contact lens use (7–9). Water for hand washing is often scarce in places where *Acanthamoeba* is endemic (e.g. Middle East, South Asia, and Northern Africa) (10). However, we recently showed that hand sanitizers kill *A. castellanii* trophozoites and cysts, providing a possible route to reducing infection (11).

*Acanthamoebae* have been used to study bacterial endosymbiosis, megaviruses, actin function in non-muscle cells, horizontal gene transfer (HGT), and host-parasite interactions. *Acanthamoebae* host pathogenic bacteria that cause pneumonia (*Legionella*), diarrhea (*Vibrio* and *Campylobacter*), or disseminated disease (*Listeria*) (12–14). *Acanthamoebae* also contain enormous double-stranded DNA viruses, which can cause respiratory infections (15). *Acanthamoeba* was used to discover and characterize roles of actin and associated proteins in the cytoskeleton, phagocytosis, and cell motility (16). Whole genome sequences of *A. castellanii* identified >500 genes derived from bacteria by HGT, which is the greatest number of any eukaryote described to date (17). A mannose-binding protein on the surface of trophozoites, secreted proteases, and pore-forming peptides each appear to contribute to contact-mediated cytolysis of corneal epithelial cells by *Acanthamoebae* (18–20).

The *Acanthamoeba* cyst wall, which forms when trophozoites are starved, is an important virulence factor, because it makes cysts resistant to surface disinfectants, sterilizing agents in contact lens solutions, and antibiotics applied to the eye (21, 22). Fifty years ago, the cyst wall was shown to contain cellulose and have two microfibril-dense layers (outer ectocyst and an inner endocyst), as well as conical structures (ostioles) that connect the layers (23, 24). During excystation, the trophozoites escape through one of the ostioles, which were not counted (25). Although the electron micrographs were beautiful, none of the cyst wall proteins were identified, and there was minimal information as to how the two layers and ostioles are formed. Further, monoclonal antibodies, potentially useful as diagnostic reagents, were made to trophozoites but not to cysts (26, 27).

To better understand its structure, we recently identified chitin and three sets of cellulose- and chitin-binding lectins in the cyst wall of *A. castellanii Neff,* which is the best studied strain of the protist (28). These lectins are present in 12 copies (Luke), 5 copies (Jonah), or 14 copies (Leo) in the cyst wall, which make them by far the most abundant proteins. These lectins contain carbohydrate-binding modules that were shown to bind cellulose in *Dictyostelium* and plants (CBM49s of Luke lectins), were previously identified in bacteria but were uncharacterized (choice of anchor A (CAA) domains of Jonah lectins), or are unique to *Acanthamoebae* (8-Cys domains of Leo lectins) (Fig. 1) (29–31). A representative Jonah lectin is present in the ectocyst layer, while representative Luke and Leo lectins are present in the endocyst layer and ostioles.

**FIG 1.**
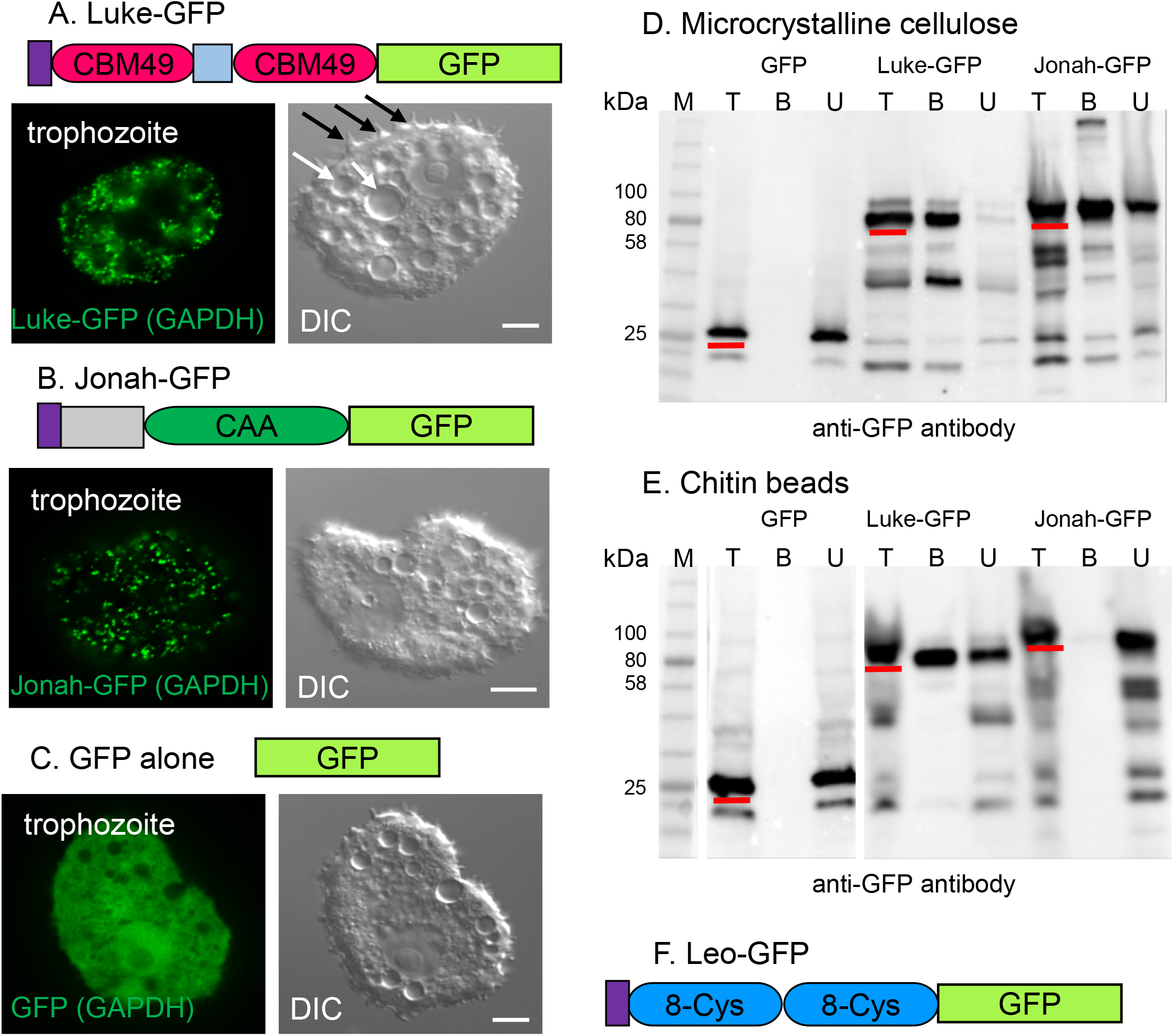
Western blots showed both Luke and Jonah lectins, which were expressed under the GAPDH promoter in small vesicles of transfected trophozoites, bound to microcrystalline cellulose, while Luke also bound to chitin beads. A. A representative Luke lectin, comprised of a signal peptide (purple) and two CBM49s separated by a Ser- and Pro-rich spacer (light blue), was tagged with GFP and expressed under a constitutive GAPDH promoter in transfected trophozoites. Luke-GFP (green) localized to small vesicles, which were distinct from larger vacuoles (white arrows) in a trophozoite that retained acanthopods on its surface (black arrows). B. Jonah-GFP, which was comprised of a signal peptide, Thr-, Lys, and Cys-rich spacer (gray), a CAA domain, and a GFP-tag, also localized to small vesicles that were distinct from larger vacuoles when expressed under the GAPDH promoter. C. In contrast, GFP alone, which was also expressed under the GAPDH promoter, diffusely labeled the cytosol, while vacuoles appeared black. D and E. Luke-GFP, Jonah-GFP, and GFP alone were released from lysed trophozoites and incubated with microcrystalline cellulose or chitin beads. Total proteins (T), bound proteins (B), and unbound proteins (U), as well as molecular weight markers (M), were run on SDS-PAGE, transferred to PVDF membranes, and detected with an anti-GFP reagent. Full-length products in total fractions are underlined in red. Luke-GFP, which included some breakdown products, bound completely to microcrystalline cellulose and partially to chitin. Jonah-GFP, which also included some breakdown products, bound partially to microcrystalline cellulose but not at all to chitin. GFP alone (negative control) did not bind to cellulose or chitin. F. Leo-GFP, which was comprised of a signal peptide, two 8-Cys domains, and a GFP-tag, was not well-expressed by trophozoites under a GAPDH promoter. A to C. Scale bars are 5 μm.

Here we used structured illumination microscopy (SIM), probes for glycopolymers, and green fluorescent protein (GFP) tags for cyst wall lectins to determine the sequence of events (stages) involved in the formation of the *A. castellanii* cyst wall during encystation (32). To characterize their carbohydrate-binding specificity, we isolated GFP-tagged lectins, which were expressed under a constitutive glyceraldehyde 3-phosphate dehydrogenase (GAPDH) promoter, from lysed trophozoites and incubated them with microcrystalline cellulose and chitin beads (33, 34). We used GFP-tagged lectins expressed under their own promoter to localize each protein in encysting protists and in mature cyst walls. To test the effect of timing of expression on localization, we compared the locations of GFP-tagged lectins expressed under their own or GAPDH promoters. We used anti-GFP antibodies to test the accessibility of GFP-tagged lectins in the ectocyst and endocyst layers of mature cyst walls. Finally, we used maltose-binding protein (MBP) fusions with Jonah, Luke, and Leo lectins to test the accessibility of glycopolymers in developing and mature cyst walls (35).

## Results

### GFP-tagged Luke and Jonah lectins each bound cellulose, while Luke lectin also bound chitin

Under their own promoters, GFP-tagged Luke, Jonah, and Leo lectins are not expressed by trophozoites but are strongly expressed in encysting protists (28) (also see below). In contrast, under a constitutive GAPDH promoter, Luke-GFP with two CBM49s and Jonah-GFP with a single CAA domain were each present in vesicles, but not on the surface, of the vast majority of transfected trophozoites (Figs. 1A and 1B). Transfected trophozoites remained motile and did not change their “acanthamoebo¡d” appearance when observed by DIC microscopy. GFP alone (negative control) was present in the cytosol of transfected trophozoites (Fig. 1C). Western blots showed Luke-GFP and Jonah-GFP, which were released from lysed trophozoites, bound to microcrystalline cellulose, while GFP alone did not bind (Fig. 1D). Although the full-length product was the major band on SDS-PAGE, both Luke-GFP and Jonah-GFP showed some degradation products. Luke-GFP also bound well to chitin, while Jonah-GFP and GFP alone did not bind to chitin (Fig. 1E). For reasons that are unclear, Leo-GFP, which has two unique 8-Cys domains, expressed weakly, if at all, under the GAPDH promoter in trophozoites, so we were unable to perform Western blots or examine mature cyst with SIM (Fig. 1F).

These results suggested that GFP-tagged lectins, expressed under the GAPDH promoter (here) or under their own promoters (next section), may serve as internal probes for cellulose +/− chitin. Overexpression of GFP-tagged Luke and Jonah lectins under the GAPDH promoter did not trigger trophozoites to encyst, while Luke and Leo lectins failed to bind to the surface of trophozoites in the absence of glycopolymers.

### In the first stage of encystation (3 to 9 hr), cellulose, chitin, and a Jonah lectin were each made in vesicles

The development of the cyst wall of *A. castellannii* was examined with external probes that bind to chitin (wheat germ agglutinin) (WGA) and to β-1,3- and β-1,4-linked polysaccharides (the fluorescent brightener calcofluor white) (CFW) in the walls of *A. castellanii, Saccharomyces,* and *Entamoeba* (28, 36–38). Glycopolymers were also visualized using a glutathione-S-transferase (GST) fusion-protein, which contains a CBM49 of a Luke lectin (28–31, 39). We recently used SIM and these probes to localize the ectocyst layer (GST-CBM49), endocyst layer (CFW), and ostioles (WGA) of mature cyst walls (28).

As early as 3 hr after placement on non-nutrient agar, Jonah-GFP expressed under its own promoter, was present in dozens of small vesicles of encysting organisms (Fig. 2A). Chitin, detected with WGA, was also made early and was present in vesicles of varying sizes, which did not overlap with those containing Jonah-GFP. In addition, the glycopolymer labeled with GST-CBM49 (most likely cellulose) was made early in dozens of small vesicles, which were distinct from those labeled with WGA (Figs. 2B and 2C). Glycopolymers labeled with CFW were not visible in organisms encysting for 3 and 6 hr, but CFW labeled a thin, spherical wall after 9 hr encystation. At this time, rare organisms had one or two small, flat ostioles, but most had none. Finally, neither Luke-GFP nor Leo-GFP, each expressed under its own promoter, was visible during this first stage of development of the cyst wall.

**FIG 2.**
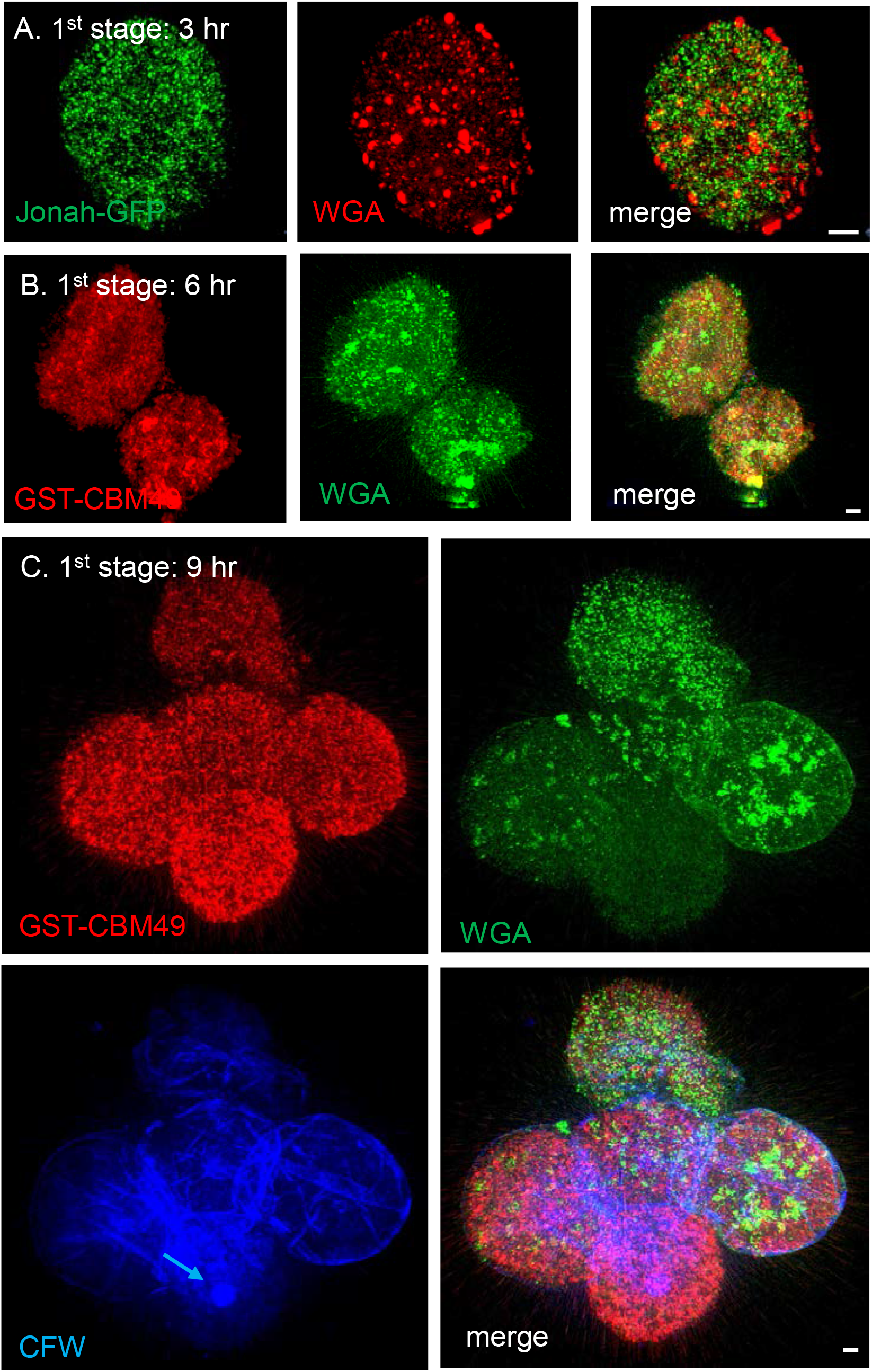
During the first stage of encystation, SIM showed Jonah-GFP, as well as glycopolymers labeled by GST-CBM49 and WGA, were present in dozens of distinct vesicles. A. After 3 hr encystation, Jonah-GFP (green), which was expressed under its own promoter, was present in dozens of small vesicles. WGA (red), which was a probe for chitin, was present in fewer but larger vesicles that did not overlap with those containing Jonah-GFP (see merge). B. After 6 hr encystation, glycopolymers labeled by GST-CBM49 (red), which was a probe for cellulose, were present in dozens of vesicles that did not overlap with those labeled by WGA (green). C. After 9 hr encystation, glycopolymers labeled with GST-CBM49 and WGA were again very abundant and present in vesicles that did not overlap. CFW, which binds to β-1,3- and β-1,4-linked polysaccharides, was not visible in vesicles of organisms encysting for 3 to 6 hr, but CFW (blue) labeled the surface of encysting protists at 9 hr. A single ostiole, which was small and circular (blue arrow), was present on the surface of one encysting protist, while ostioles were absent from the other organisms. When expressed under its own promoter, neither Luke-GFP nor Leo-GFP was present in the first stage of encystation. A to C. Scale bars are 2 μm.

These results showed that the first stage of encystation is an abrupt event in which amoeboid trophozoites rapidly synthesize glycopolymers and a Jonah lectin in dozens of vesicles that fill the encysting cells. In contrast, Luke and Leo lectins were not yet made, suggesting encystation-specific proteins are expressed at different times (40).

### In the second stage of encystation (12 to 18 hr), a primordial cyst wall was made that contains small flat ostioles and three lectins in distinct distributions

GST-CBM49, WGA, and CFW each labeled primordial cyst walls, which had a single, thin layer and small, flat ostioles (Fig. 3A). Ostioles, which labeled with CFW but not with GST-CBM49 or WGA, were at first filled circles but later became rings (Fig. 3B). While it was difficult to count these small ostioles because of variable labeling with CFW, they appeared to be in similar number (~9 per cell) and in a similar distribution as conical ostioles of mature cyst walls (see below).

**FIG 3.**
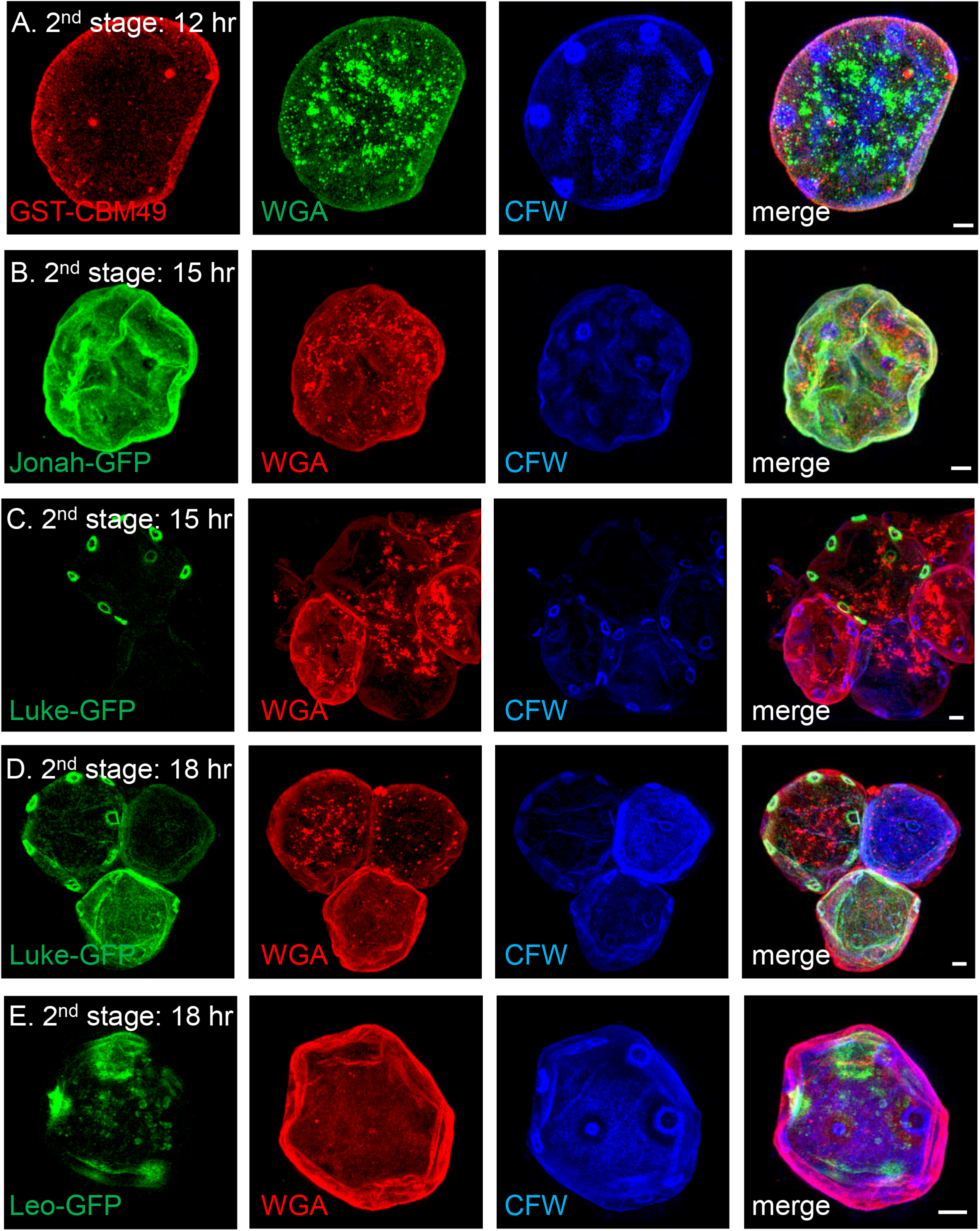
During the second stage of encystation, SIM showed a primordial cyst wall contained Jonah-GFP and glycopolymers labeled with GST-CBM49 and WGA, each in a diffuse pattern, while small, flat ostioles contained Luke-GFP and labeled with CFW. A. After 12 hr encystation, GST-CBM49 (red) diffusely labeled a thin, primordial wall, which contained small, flat ostioles visible only with CFW (blue). WGA (green), which predominantly labeled vesicles, also labeled the primordial wall. After 15 hr encystation, Jonah-GFP (green), expressed under its own promoter, was homogenously distributed in the primordial wall (B), while Luke-GFP (green), also expressed under its own promoter, outlined the edges of small ostioles in some cells (C). After 18 hr encystation, Luke-GFP, which continued to outline the edges of small ostioles, also spread across the surface of some primordial walls (D), while Leo-GFP, expressed under its own promoter, was in a patchy distribution in primordial walls that was, for the most part, independent of ostioles (E). A to E. Scale bars are 2 μm.

Each of the GFP-tagged lectins expressed under its own promoter was present in primordial ectocyst walls but in markedly different distributions. Jonah-GFP was homogenously distributed across the surface of the primordial ectocyst wall (Fig. 3B). Luke-GFP outlined some but not all of early ring-shaped ostioles (Fig. 3C). Later, in addition to outlining the ostioles, Luke-GFP was homogenously distributed across the surface of the primordial cyst wall (Fig. 3D). Leo-GFP was latest to the wall and had a patchy distribution, which was, for the most part, independent of the ostioles (Fig. 3E).

In summary, the primordial cyst walls contained the glycopolymers and lectins present in mature cyst walls but in a single thin layer. The presence of small, circular ostioles, which were visualized by the external probe CFW or the internal probe Luke-GFP, showed these structures are initiated prior to separation of the ectocyst and endocyst layers.

### In the third stage of encystation (24 to 36 hr), the ectocyst and endocyst layers separated and the ostioles became conical

In the third stage and in mature cyst walls (≥72 hr), the cell body contracted, so that the emerging endocyst layer was made inside the ectocyst layer (Figs. 4 and 5). Glycopolymers labeled by CFW moved to the endocyst layer and ostioles, which were often labeled with WGA, the probe for chitin. Jonah-GFP remained with the ectocyst layer and had essentially the same appearance in the walls of second stage, third stage, and mature cysts (Figs. 3B, 4A, 4D, and 5A). Luke-GFP, which was diffusely distributed in the endocyst layer and dome-shaped ostioles of organisms encysting for 24 and 36 hr, sharply outlined conical ostioles in mature cyst walls (Figs. 4B, 4E, and 5C). For the most part, Leo-GFP localized the endocyst layer of 24 hr cysts, although its distribution remained patchy (Fig. 4C). It was not until 36 hr encystation that Leo-GFP began to diffusely label the endocyst layer and outline ostioles, which was its distribution in mature cysts (Figs. 4F and 5E).

**FIG 4.**
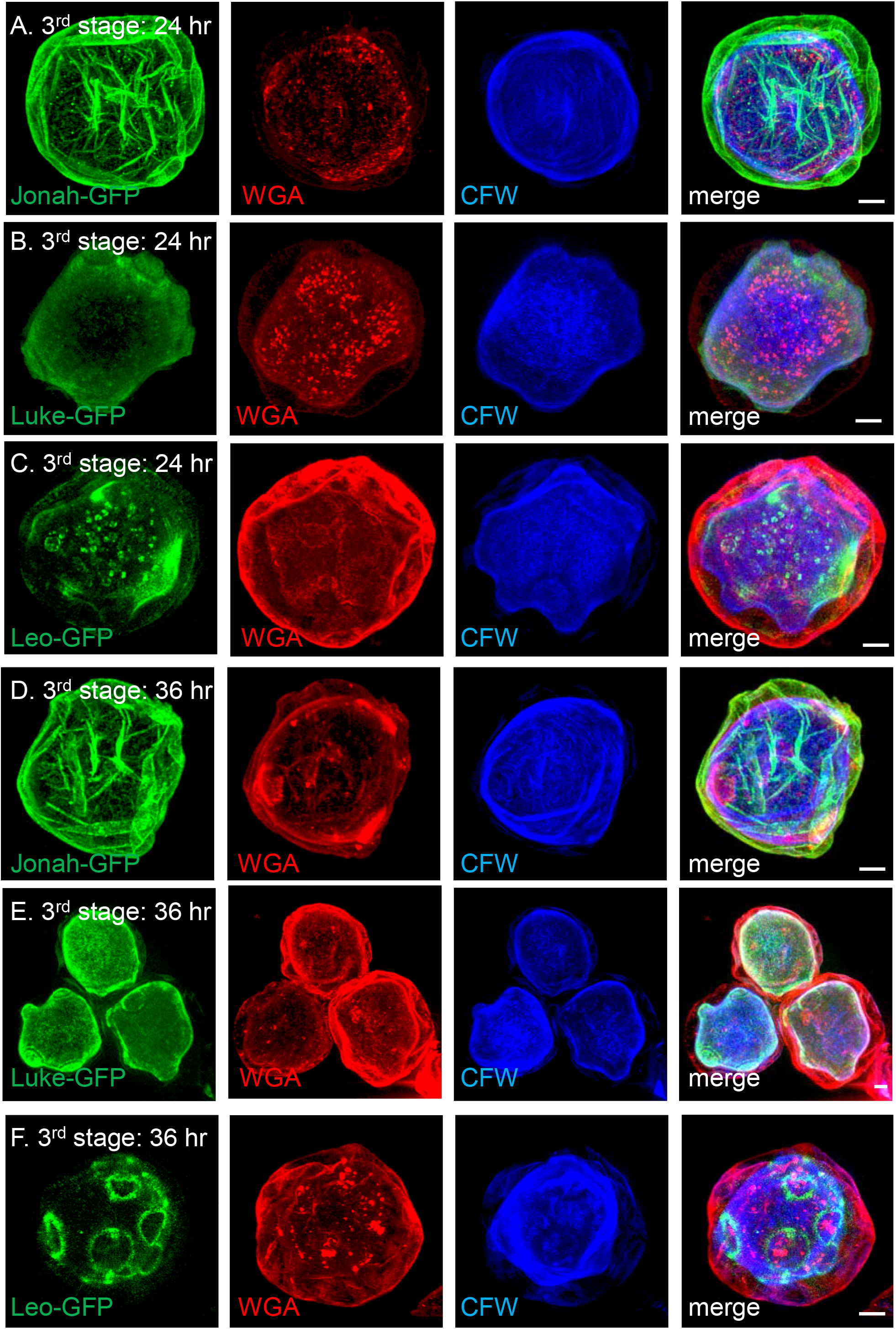
During the third stage of encystation, SIM showed Jonah-GFP remained in the ectocyst layer, while Luke-GFP and Leo-GFP moved to the endocyst layer and ostioles. Each of the GFP-tagged lectins was expressed under its own promoter. Jonah-GFP (green) was abundant in the ectocyst layer of walls of protists encysting for 24 hr (A) or 36 hr (D). Luke-GFP was homogeneously distributed in the endocyst layer, as well as cone-shaped ostioles, at 24 hr (B) and 36 hr (E) encystation. Leo-GFP was present in vesicles and in a somewhat patchy distribution in both the ectocyst and endocyst layers of organisms encysting for 24 hr (C). It was not until 36 hr (F) that Leo-GFP began to sharply outline ostioles. CFW (blue) labeled the endocyst layer and ostioles, while WGA (red) labeled the endocyst layer (A and B), ectocyst layer (C and F), or both layers (D and E). A to E. Scale bars are 2 μm.

**FIG 5.**
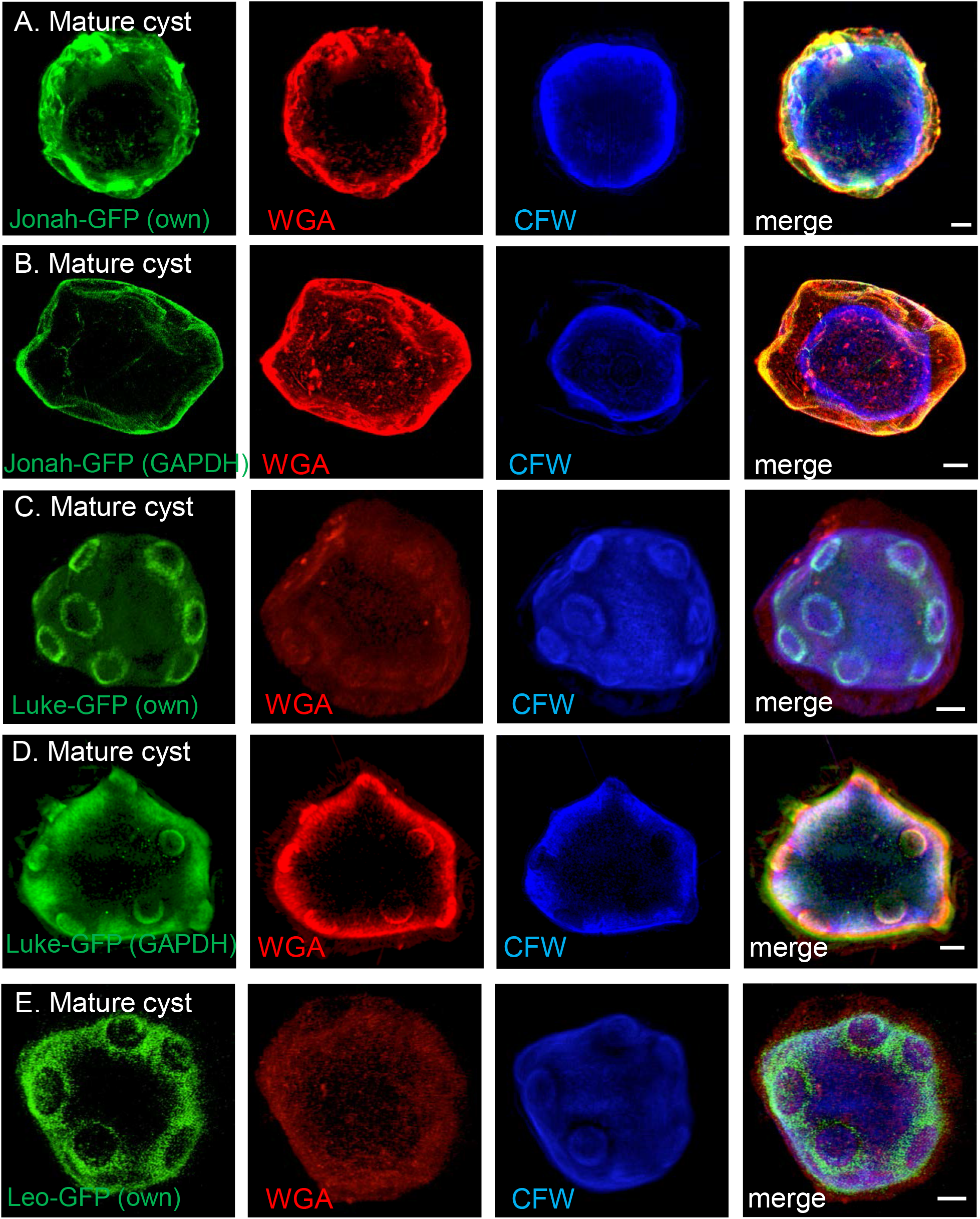
SIM showed GFP-tagged Jonah and Luke lectins localized to the same places in mature cyst walls when each was expressed under its own encystation-specific promoter or under a constitutive GAPDH promoter. Jonah-GFP (green) expressed under its own promoter (A) or the GAPDH promoter (B) localized to the ectocyst layer, which was also labeled with WGA (red). The endocyst layer was labeled blue with CFW. Luke-GFP labeled the endocyst layer and outlined the ostioles when expressed under its own promoter (C) or the GAPDH promoter (D). Leo-GFP, which expressed well under its own promoter (E) but was not expressed under the GAPDH promoter, also labeled the endocyst layer and outlined the ostioles. A to E. Scale bars are 2 μm.

In summary, while Jonah-GFP arrived early at its final destination in the ectocyst layer, Luke and Leo lectins, by somewhat different routes, arrived much later to their final destinations in the endocyst layer and ostioles. In a similar way, the ostioles, which were sharply outlined by Luke and Leo lectins, did not obtain their distinctive conical shape until late in encystation (between 36 and 72 hr).

### Timing of expression of GFP-tagged lectins did not affect their locations in mature cyst walls

Jonah-GFP expressed under a constitutive GAPDH promoter had the same location in the ectocyst layer of mature walls, as did Jonah-GFP expressed under its own promoter (Figs. 5A and 5B). Luke-GFP expressed under either the GAPDH promoter or its own promoter had the same locations in endocyst layer and ostioles of mature walls (Figs. 5C and 5D). Because Leo-GFP did not express well under the GAPDH promoter, it was not possible to compare its distribution versus Leo-GFP under its own promoter (Fig. 5E). GFP alone expressed under the GAPDH promoter (negative control) remained in the cytosol of cysts (data not shown). These results suggested carbohydrate-binding specificities or protein-protein interactions were more important than timing of expression for localization of Jonah and Luke lectins.

### GFP-tagged lectins and glycopolymers were accessible in ectocyst layer but were inaccessible in the endocyst layer and ostioles of mature cyst walls

To determine the accessibility of proteins in the ectocyst and endocyst layers and ostioles of mature cyst walls, we incubated organisms expressing GFP-tagged lectins under their own promoters with anti-GFP antibodies. Widefield microscopy showed that Jonah-GFP was accessible in the endocyst layer of nearly 100% of cysts with a detectable Jonah-GFP signal (Fig. 6A). In contrast, anti-GFP antibodies showed Luke-GFP and Leo-GFP were accessible in the endocyst layer and ostioles of 3 and 2%, respectively, of cysts with detectable GFP signals (Figs. 6B and 6C).

**FIG 6.**
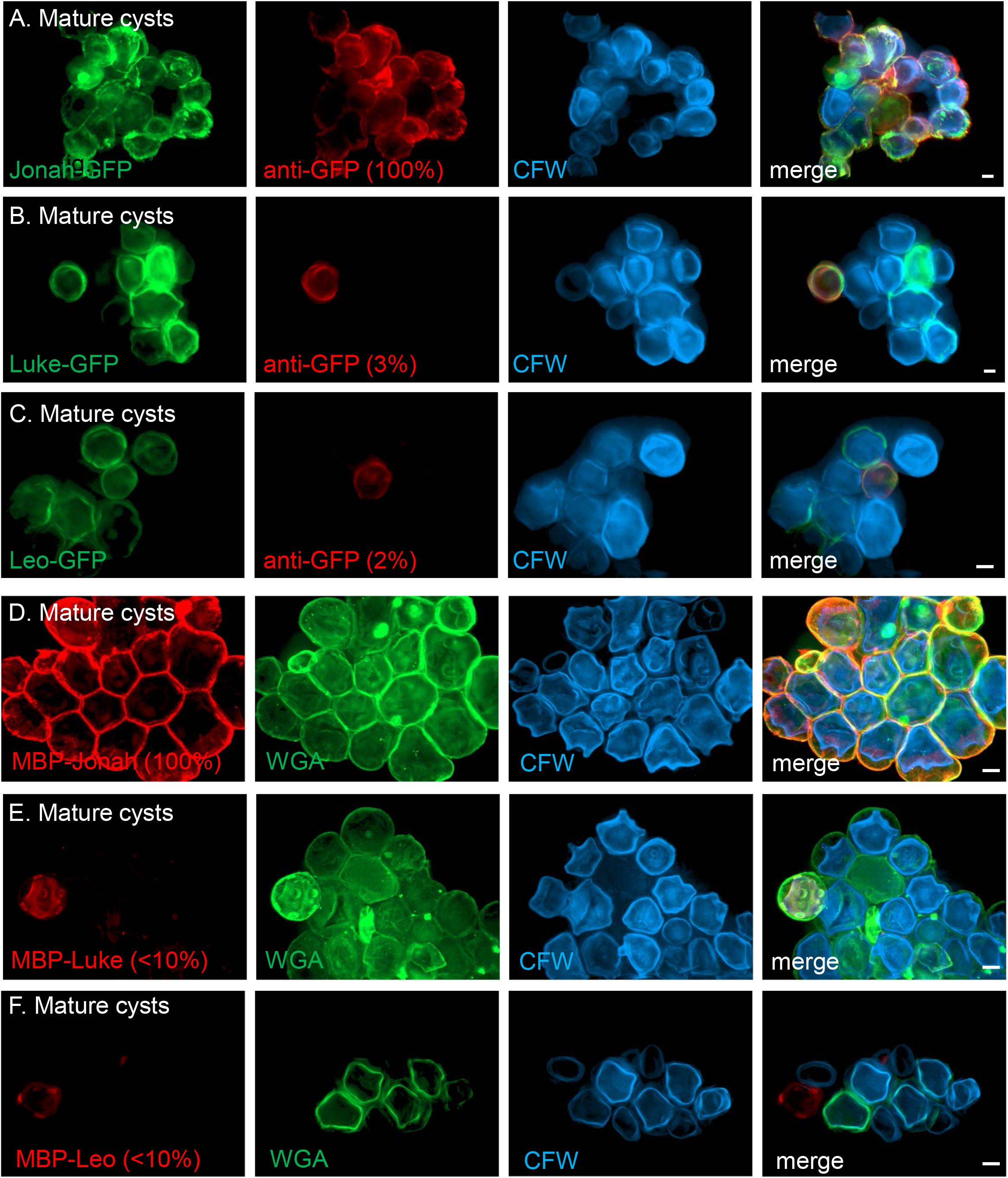
Widefield microscopy showed Jonah-GFP and glycopolymers labeled by MBP-Jonah were accessible in the ectocyst layer of mature cyst walls, while Luke-GFP, Leo-GFP, and glycopolymers labeled by MBP-Luke and MBP-Leo were inaccessible in the endocyst layer and ostioles. A. Nearly 100% of mature cysts expressing Jonah-GFP (green) under its own promoter were labeled with anti-GFP antibodies (red). CFW (blue) labeled the endocyst layer of cyst walls. B. Just 3% of mature cysts expressing Luke-GFP under its own promoter labeled with anti-GFP antibodies. C. Just 2% of mature cysts expressing Leo-GFP under its own promoter labeled with anti-GFP antibodies. D. MBP-Jonah (red) labeled 100% of mature cysts, which were also labeled with WGA (green) and CFW. In contrast, MBP-Luke (E) and MBP-Leo (F) each labeled <10% of mature cysts. A to F. Scale bars are 5 μm. SIM of mature cysts labeled with MBP-lectin fusions are shown in Figs. 7D to 7F.

To determine the accessibility of glycopolymers in primordial and mature cyst walls, we labeled both sets of organisms with MBP-Jonah, MBP-Luke, and MBP-Leo, each of which was made as recombinant protein in the periplasm of *E. coli.* We recently showed all three MBP-fusions bind cellulose, while binding to chitin was more variable (28). Each MBP-fusion bound to glycopolymers in vesicles and in primordial walls of 100% of second stage organisms (Figs. 7A to 7C). MBP-Jonah also bound to the ectocyst layer of 100% of mature cyst walls (Figs. 6D and 7D), which was the same location as Jonah-GFP expressed under either its own or the GAPDH promoter (Figs. 5A and 5B). In contrast, MBP-Luke and MBP-Leo probes labeled the endocyst layer and ostioles of <10% mature cyst walls (Figs. 6E, 6F, 7E, and 7F). Although these were the same places in mature cyst walls where Luke-GFP and Leo-GFP localized under their own promoters or the GAPDH promoter (Luke-GFP) (Figs. 5C to 5E), these results suggested that glycopolymers bound by Luke and Leo lectins in the endocyst layer and ostioles were, for the most part, inaccessible to external probes.

**FIG 7.**
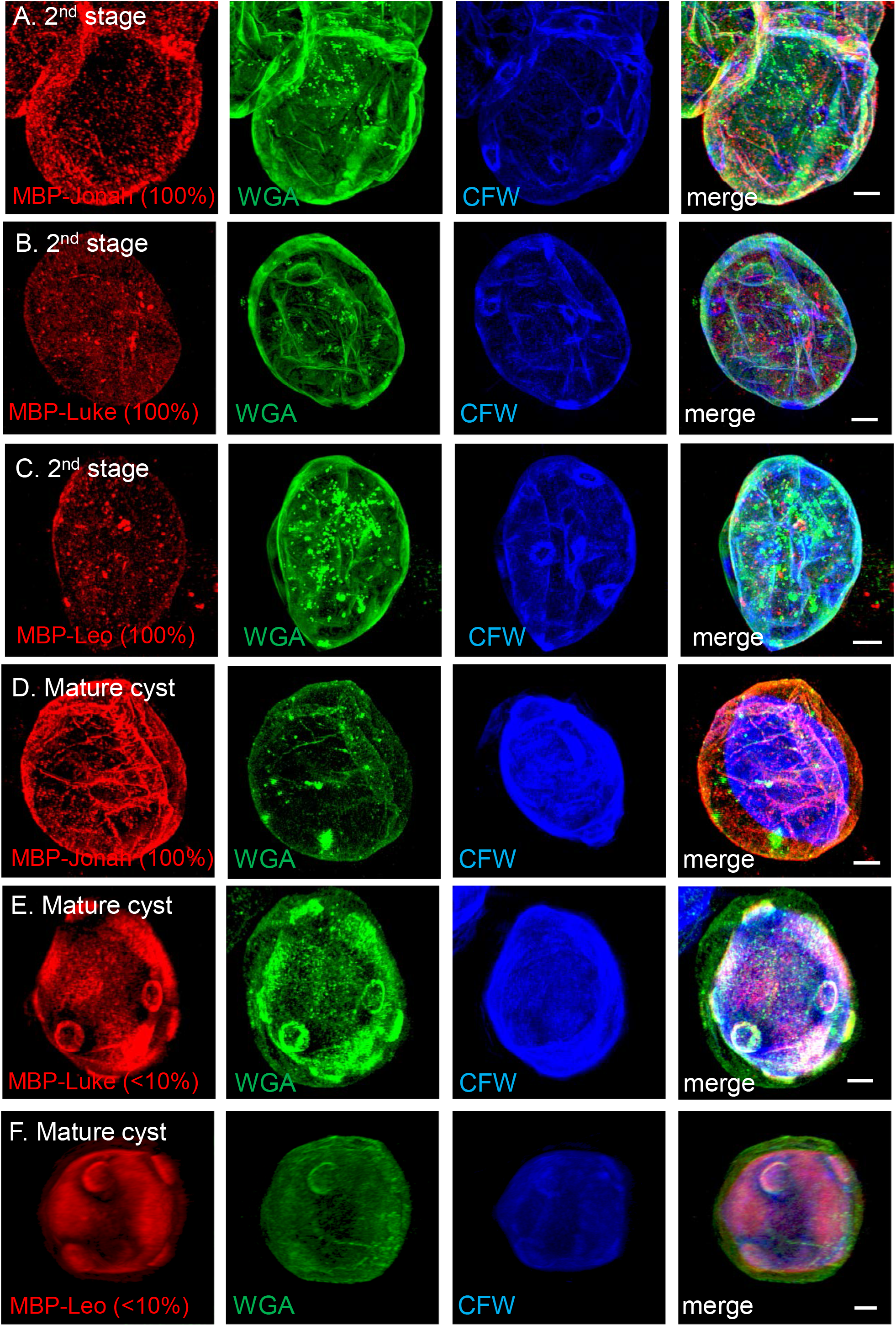
SIM showed glycopolymers bound by MBP-lectin fusions were accessible in the primordial wall and in the ectocyst wall but were inaccessible in the endocyst wall and ostioles. After 12 hr encystation (second stage), MBP-Jonah (A), MBP-Luke (B), and MBP-Leo (C) labeled vesicles and primordial walls in nearly 100% of organisms. WGA diffusely labeled primordial walls, while CFW highlighted ring-shaped ostioles. Although MBP-Jonah labeled the ectocyst layer of nearly 100% of mature cysts (D), MBP-Luke (E) and MBP-Leo (F) each labeled the endocyst layer and ostioles of <10% of mature cysts. A to F. Scale bars are 2 μm. Widefield micrographs of groups of mature cysts labeled with MBP-lectin fusions are shown in Figs. 6D to 6F.

Finally, by rotating three-dimensional SIM reconstructions of organisms expressing Luke-GFP or Leo-GFP or labeled with WGA, MBP-Luke, or MBP-Leo, we counted an average of 8.8 +/− 2.5 ostioles per mature cyst wall (24 cysts total). To our knowledge, this is the first estimate of the number of ostioles in *Acanthamoeba* cyst walls, because ostioles have not previously been visualized by light microscopy and are extremely difficult to count by TEM, unless dozens of serial sections are examined.

## Discussion

While our recent study provided a parts list of *A. castellanii* cyst wall proteins (Jonah, Luke, and Leo lectins) and glycopolymers (cellulose and chitin) (28), the present study identified the major events involved in making the two layers of the cyst wall and ostioles. While the boundaries between the three stages of the development of the cyst wall were somewhat arbitrary (based upon the times selected for examining cyst walls with SIM), each stage had several distinguishing features.

In the first stage, encysting organisms rapidly transformed from amoeboid trophozoites, which were full of vacuoles and had acanthopods, to immotile, synthetic forms making glycopolymers and Jonah lectins in dozens of distinct vesicles. Encysting *Entamoebae* also transform from amoeboid trophozoites to immotile, synthetic forms making chitin and the Jacob lectin in distinct vesicles (41). In an analogous way, encysting *Giardia* transform from flagellated forms with an adherence disc to spherical, immotile forms making β-1,3-linked GalNAc glycopolymer and cyst wall proteins (CWP1 to CWP3) in distinct vesicles (42). In contrast, no dramatic secretory event occurs in fungi or plants, which remodel their walls with growth, differentiation, or cell division, but never make their walls from scratch, with exception of the septum separating dividing cells (43, 44).

Because vesicles labeled with WGA and GST-CBM49 did not overlap, it is likely that chitin and cellulose synthases are present in distinct compartments. Definitive identification of cellulose and chitin in vesicles of encysting organisms will depend upon localization of tagged cellulose and chitin synthases, each of which is encoded by a single gene in *A. castellanii* (17, 28, 30, 45, 46). Further, expression of tagged synthases under a GAPDH promoter in trophozoites will determine whether accessory proteins are necessary for making cellulose or chitin. In yeast, three chitin synthases (Chs1 to Chs3) and four auxiliary proteins (Chs4 to Chs7) are precisely localized in the plasma membrane by actin and ras homolog proteins such as RhoA and Cdc42 (43, 47, 48). In plant cells, three cellulose synthases form a plasma membrane complex, which is localized by cortical microtubules in association with Rabs, SNAREs, and other transport-associated proteins (49–51). While *A. castellanii* has homologs of most of the yeast and plant proteins involved in localizing chitin synthases in fungi and cellulose synthases in plants, examination of their roles in chitin and cellulose synthesis was beyond the scope of the present study (45, 46).

In the second stage, the two most remarkable features of the primordial cyst wall were the distinct distributions of the GFP-tagged lectins and the sets of small, circular ostioles. Early on, Jonah-GFP was homogenously distributed across the primordial wall, while Luke-GFP outlined some ostioles. Later, Luke-GFP spread across the primordial wall, while Leo-GFP appeared in patches, which were not specific to any structure. Indeed, it appeared that Luke-GFP was secreted onto the surface of second stage organisms but did not find the glycopolymer to which it binds.

Because the ostioles labeled with the least specific external probe (CFW) but not with GST-CBM49 or WGA, it was not possible to determine whether ostioles are composed of cellulose, chitin, or another glycopolymer. Indeed CFW has been shown to bind numerous β-1,3 and β-1,4 polysaccharides, including cellulose, chitin, mixed linkage glucans, and galactoglucomannan (52). The presence of the internal probe Luke-GFP in the small ostioles did not settle this problem, as Luke-GFP released from lysed trophozoites bound to both microcrystalline cellulose and chitin beads. As ostioles labeled with CFW before they contained Luke-GFP, it is likely that glycopolymers are the drivers behind the circular or ring-like structures. How the small ostioles simultaneously appear in their eventual distribution on the surface of the primordial cyst wall and synchronously develop into conical structures was not determined here. Because *A. castellanii* has almost nine ostioles but uses just one for the excysting trophozoite to escape the cyst wall, it is likely that ostioles serve other functions such as holding layers of the cyst wall together and/or exchanging nutrients or waste products with the environment. How *A. castellanii* glycoside hydrolases, which belong to multiple CAZy families (GH1, GH5, GH8, GH10, GH16, and GH18), are involved in wall formation during encystation or enzyme-catalyzed destruction of the ostioles during excystation is also of great interest but beyond the scope of these studies (30, 45, 46).

In the third stage, the Jonah lectin and the glycopolymer bound by Jonah remained in the ectocyst layer, while Luke and Leo lectins and the glycopolymers bound by the lectins moved to the endocyst layer and ostioles. In the same way, the outer primary layer of plant cells forms before the inner secondary layer (53). The distinct distributions of the three GFP-tagged lectins in the primordial, third stage, and mature cyst walls strongly suggests each lectin binds to different glycopolymers (e.g. cellulose versus chitin), glycopolymers modified in different ways (e.g. unmodified chitin versus deacetylated chitosan), and/or glycopolymers with different microfibrillar structures (e.g. microcrystalline versus amorphous cellulose). In support of this idea, each lectin had the same localization in mature cyst walls when expressed as an internal probe with a GFP-tag under its own or under a constitutive GAPDH promoter or when applied externally as an MBP-fusion. There may also be protein-protein interactions and/or lectin-glycoprotein interactions, which determine the localization of cyst wall lectins in the *A. castellanii* cyst wall. As an example of protein-protein interactions, an *E. histolytica* Jessie lectin has a chitin-binding domain and a self-agglutinating “daub” domain, which makes cyst walls impermeable to small probes such as phalloidin (41). As an example of lectin-glycan interactions, the Gal/GalNAc lectin on the surface of *Entamoebae* binds to glycans on Jacob lectins, which, in turn, bind to chitin fibrils in the cyst wall (38).

Jonah-GFP and glycopolymers bound by MBP-Jonah were accessible in the ectocyst layer of mature cyst walls, while Luke-GFP and Leo-GFP and glycopolymers bound by MBP-Luke and MBP-Leo were, for the most part, inaccessible in the endocyst layer and ostioles. It appears then that cyst wall lectins and glycopolymers in the ectocyst layer block access of external probes to the endocyst layer and ostioles. Alternatively, cyst wall lectins and glycopolymers in the endocyst layer and ostioles are so tightly packed that they are inaccessible to external probes. Because of its abundance and accessibility in the ectocyst layer, the Jonah lectin appears to be an excellent target for diagnostic antibodies for *A. castellanii* cysts. We expect each antibody would react with a single Jonah lectin, as paralogous Jonah lectins share less than 40% amino acid identities (28). In contrast, Luke and Leo lectins were inaccessible in the endocyst layer and ostioles, and so these abundant cyst wall proteins do not appear to be good targets for diagnostic anti-cyst antibodies.

## MATERIALS AND METHODS

### Culture of trophozoites and preparation of cysts

Culture and manipulation of *Acanthamoebae* have been approved by the Boston University Institutional Biosafety Committee. *A. castellanii* Neff strain trophozoites, which included the parent strain and transfectants expressing GFP-tagged cyst wall proteins (see next section), were cultured in ATCC 712 medium at 30°C and induced to encyst on non-nutrient agar plates, again at 30°C, using methods that have recently been described (28, 54, 55). After 3, 6, 9, 12, 15, 18, 24, 36, or ≥72 hr incubation, 15 ml of phosphate-buffered saline (PBS) was added to agar plates, which were incubated on a shaker for 30 min at room temperature (RT) to release organisms. Encysting organisms and cysts were all washed in PBS and then fixed in 1% paraformaldehyde buffered with phosphate for 15 min at RT. Organisms were washed with Hank’s Buffered Saline Solution (HBSS), incubated with HBSS containing 1% bovine serum albumin (BSA) for 1 hour at RT, and used immediately for labeling and SIM or stored at 4^0^C.

### Expression of GFP-tagged cyst wall proteins in transfected *A. castellanii*

Previously, we made stable *A. castellanii* transfectants, which expressed Jonah lectin (ACA1_164810), Luke lectin (ACA1_377670), or Leo lectin (ACA1_074730), each with a C-terminal GFP-tag and under its own promoter (~500-bp of the 5’ UTR) (28, 33, 34). Here we also made stable transfectants, which expressed Jonah-GFP and Luke-GFP under a constitutive (GAPDH) promoter. The methods for transfecting trophozoites with SuperFect Transfection Reagent and for selecting transfectants with G418 were exactly as described previously (28). After 2 to 4 weeks antibiotic selection, trophozoites expressing GFP-fusions under the GAPDH promoter were fixed with paraformaldehyde and observed with widefield and differential interference contrast microscopy, using a 100x objective of a Zeiss AXIO inverted microscope with a Colibri LED (Carl Zeiss Microcopy LLC, Thornwood, NY), as described previously (28).

### Binding of GFP-tagged lectins to commercially available glycopolymers

To test the carbohydrate-binding specificity of the GFP-tagged lectins, we lysed trophozoites expressing each protein under a GAPDH promoter and then incubated the lysate with microcrystalline cellulose or chitin beads, using methods to characterize Jonah, Luke, and Leo lectins fused to maltose-binding protein (MBP) (28, 29, 35). Total (T) proteins prior to incubation with glycopolymers, unbound proteins (U), and bound proteins (B) released with SDS were separated on SDS-PAGE, transferred to PVDF, and detected with reagents that recognize GFP. A control was GFP alone, which was expressed under a GAPDH promoter in the cytosol of transfected *A. castellanii* trophozoites.

### External probes for labeling encysting organisms and SIM methods

A GST-CBM49 fusion-construct, which contains the N-terminal CBM49 of a Luke lectin minus its signal peptide, was made in the cytosol of bacteria, purified on glutathione-agarose, and labeled with Alexafluor 594, as recently described (28, 29, 39). After fixation with formaldehyde and blocking with BSA (above), non-transfected organisms encysting for zero to 12 hr were incubated for 2 hr at 4°C with 2.5 μg GST-CBM49 conjugated to Alexa Fluor 594 and 12.5 μg of WGA (Vector Laboratories, Burlingame, CA) conjugated to Alexa Fluor 488 in 100 μl HBSS. Alternatively, transfectants expressing Jonah-GFP, Luke-GFP, or Leo-GFP were encysted for zero to ≥72 hr, fixed, blocked, and then labeled with WGA conjugated to Alexa Fluor 594. Samples were then labeled with 100 μg of calcofluor white M2R (Sigma-Aldrich) in 100 μl HBSS for 15 min at RT and washed five times with HBSS. Preparations were observed with widefield and differential interference contrast microscopy, as described above. Alternatively, preparations were mounted in Mowiol mounting medium (Sigma-Aldrich), and SIM was performed with a 63-x objective of a Zeiss ELYRA S.1 microscope at Boston College (Chestnut Hill, MA), and 0.09-μm optical sections deconvolved using Zen software, as previously described (28, 32).

We used anti-GFP antibodies to determine the accessibility of GFP-tagged lectins in mature cyst wall. Without prior fixation, mature cysts expressing GFP-fusions under their own promoter were blocked with BSA, incubated with 1:400 mouse anti-GFP IgG (Roche) for 1 hr at RT, washed, and then incubated with 1:800 goat anti-mouse IgG-Alexa Fluor 594 (Molecular Probes, Invitrogen). Preparations were labeled washed, labeled with WGA and CFW, fixed in paraformaldehyde, mounted on glass slides, and observed with widefield microscopy. To determine the accessibility of glycopolymers in primordial and mature cyst walls, we used MBP-fusions to Luke, Leo, and Jonah lectins, which were previously made to characterize the carbohydrate-binding specificity of each lectin (28). Organisms encysting for 12 hr (second stage) and ≥72 hr (mature cysts) were fixed, blocked, and incubated with 15 μg of each MBP-CWP fusion conjugated to Alex Fluor 594 for 2 hr at 4°C. Preparations were labeled with WGA conjugated to Alexa Fluor 488 and CFW, as described above, and visualized with widefield microscopy and SIM. Finally, to count the number of ostioles per cyst wall, we rotated three-dimensional SIM reconstructions of mature cysts expressing Luke-GFP or Leo-GFP or non-transfectants labeled with WGA, MBP-Luke, or MBP-Leo, all of which clearly outlined conical ostioles.

## Acknowledgments

We thank Maria Ericsson of Harvard University for help with transmission electron microscopy. We thank Bret Judson and Marc-Jan Gubbels of Boston College for help with structured illumination microscopy, which was supported by NSF grant No. 1626072. This work was also supported by awards from the National Institute of Allergy and Infectious Diseases of the NIH to J.S. (R01 AI110638), from the National Institute of General Medical Science of the NIH to C.E.C. (P41 GM104603), and from the Department of Energy to B.R.U. (DESC0015662).

## REFERENCES

1. Taylor HR, Burton MJ, Haddad D, West S, Wright H. Trachoma. Lancet 2014;384:2142–52

2. Ma LJ, Geiser DM, Proctor RH, Rooney AP, O’Donnell K, Trail F, et al. *Fusarium* pathogenomics. Annu Rev Microbiol 2013;67:399–416

3. Lorenzo-Morales J, Khan NA, Walochnik J. An update on *Acanthamoeba* keratitis: diagnosis, pathogenesis and treatment. Parasite 2015;22:10

4. Carrijo-Carvalho LC, Sant’ana VP, Foronda AS, de Freitas D, de Souza Carvalho FR. Therapeutic agents and biocides for ocular infections by free-living amoebae of *Acanthamoeba* genus. Surv Ophthalmol 2017;62:203–18

5. Carnt N, Robaei D, Minassian DC, Dart JKG. *Acanthamoeba* keratitis in 194 patients: risk factors for bad outcomes and severe inflammatory complications. Br J Ophthalmol 2018;102:1431–35

6. Satlin MJ, Graham JK, Visvesvara GS, Mena H, Marks KM, Saal SD, et al. Fulminant and fatal encephalitis caused by *Acanthamoeba* in a kidney transplant recipient: case report and literature review. Transpl Infect Dis 2013;15:619–26

7. Lorenzo-Morales J, Martín-Navarro CM, López-Arencibia A, Arnalich-Montiel F, Piñero JE, Valladares B. *Acanthamoeba* keratitis: an emerging disease gathering importance worldwide? Trends Parasitol 2013;29:181–7

8. Cope JR, Collier SA, Rao MM, Chalmers R, Mitchell GL, Richdale K, et al. Contact lens wearer demographics and risk behaviors for contact lens-related eye infections--United States, 2014. MMWR Morb Mortal Wkly Rep 2015;64:865–70

9. Carnt N, Hoffman JJ, Verma S, Hau S, Radford CF, Minassian DC, et al. *Acanthamoeba* keratitis: confirmation of the UK outbreak and a prospective case-control study identifying contributing risk factors. Br J Ophthalmol 2018:312544.

10. Mekonnen MM, Hoekstra AY. Four billion people facing water scarcity. Sci Adv 2016;2:e1500323

11. Aqeel Y, Rodriguez R, Chatterjee A, Ingalls RR, Samuelson J. Killing of diverse eye pathogens *(Acanthamoeba castellanii, Fusarium solani*, and *Chlamydia trachomatis)* with alcohols. PLoS Negl Trop Dis 2017;11:e0005382

12. Tosetti N, Croxatto A, Greub G. Amoebae as a tool to isolate new bacterial species, to discover new virulence factors and to study the host-pathogen interactions. Microb Pathog 2014;77:125–30

13. Van der Henst C, Scrignari T, Maclachlan C, Blokesch M. An intracellular replication niche for *Vibrio cholerae* in the amoeba *Acanthamoeba castellanii*. ISME J 2016;10:897–910

14. Vieira A, Seddon, Karlyshev AV. *Campylobacter-Acanthamoeba* interactions. Microbiology 2015; 161:933–47

15. La Scola B. Looking at protists as a source of pathogenic viruses. Microb Pathog 2014;77:131–5

16. Ostap EM, Maupin P, Doberstein SK, Baines IC, Korn ED, Pollard TD. Dynamic localization of myosin-I to endocytic structures in *Acanthamoeba*. Cell Motil Cytoskeleton 2003;54:29–40

17. Clarke M, Lohan AJ, Liu B, Lagkouvardos I, Roy S, Zafar N, et al. Genome of *Acanthamoeba castellanii* highlights extensive lateral gene transfer and early evolution of tyrosine kinase signaling. Genome Biol 2013;14:R11

18. Garate M, Cubillos I, Marchant J, Panjwani N. Biochemical characterization and functional studies of *Acanthamoeba* mannose-binding protein. Infect Immun. 2005;73:5775–81

19. Kim WT, Kong HH, Ha YR, Hong YC, Jeong HJ, Yu HS, Chung DI. Comparison of specific activity and cytopathic effects of purified 33 kDa serine proteinase from *Acanthamoeba* strains with different degree of virulence. Korean J Parasitol. 2006;44:321–30

20. Michalek M, Sönnichsen FD, Wechselberger R, Dingley AJ, Hung CW, Kopp A, et al. Structure and function of a unique pore-forming protein from a pathogenic acanthamoeba. Nat Chem Biol 2013;9:37–42

21. Coulon C, Collignon A, McDonnell G, Thomas V. Resistance of *Acanthamoeba* cysts to disinfection treatments used in health care settings. J Clin Microbiol 2010;48:2689–97

22. Dupuy M, Berne F, Herbelin P, Binet M, Berthelot N, Rodier MH, et al. Sensitivity of free-living amoeba trophozoites and cysts to water disinfectants. Int J Hyg Environ Health 2014;217:335–9

23. Bowers B, Korn ED. The fine structure of *Acanthamoeba castellanii* (Neff strain). II. Encystment. J Cell Biol 1969;41:786–805

24. Potter JL, Weisman RA. Cellulose synthesis by extracts of *Acanthamoeba castellanii* during encystment. Stimulation of the incorporation of radioactivity from UDP-(^14^C)glucose into alkali-soluble and insoluble beta-glucans by glucose 6-phosphate and related compounds. Biochim Biophys Acta 1976;428:240–52

25. Chávez-Munguía B, Omaña-Molina M, González-Lázaro M, González-Robles A, Bonilla P, Martínez-Palomo A. Ultrastructural study of encystation and excystation in *Acanthamoeba castellanii*. J Eukaryot Microbiol 2005;52:153–8

26. Hiwatashi E, Tachibanabi H, Kanedab Y, Obazawaa H. Production and characterization of monoclonal antibodies to *Acanthamoeba castellanii* and their application for detection of pathogenic *Acanthamoeba spp*. Parasitol Internat 1997;46:197–205

27. Kang AY, Park AY, Shin HJ, Khan NA, Maciver SK, Jung SY. Production of a monoclonal antibody against a mannose-binding protein of *Acanthamoeba culbertsoni* and its localization. Exp Parasitol 2018;192:19–24

28. Magistrado-Coxen P, Aqeel Y, Lopez A, Haserick JR, Urbanowicz BR, Costello CE, Samuelson J. The most abundant cyst wall proteins of *Acanthamoeba castellanii* are three sets of lectins that bind cellulose and chitin and localize to distinct structures in cyst walls. BioRXIV:496307 doi: https://doi.org/10.1101/496307

29. Urbanowicz BR, Catalá C, Irwin D, Wilson DB, Ripoll DR, Rose JK. A tomato endo-beta-1,4-glucanase, SlCel9C1, represents a distinct subclass with a new family of carbohydrate binding modules (CBM49). J Biol Chem 2007;282:12066–74

30. Lombard V, Golaconda Ramulu H, Drula E, Coutinho PM, Henrissat B. The Carbohydrate-active enzymes database (CAZy) in 2013. Nucleic Acids Res 2014;42:D490–D495

31. Wang Y, Slade MB, Gooley AA, Atwell BJ, Williams KL. Cellulose-binding modules from extracellular matrix proteins of *Dictyostelium discoideum* stalk and sheath. Eur J Biochem 2001;268:4334–45.

32. Demmerle J, Innocent C, North AJ, Ball G, Müller M, et al. Strategic and practical guidelines for successful structured illumination microscopy. Nat Protoc 2017; 12:988–1010

33. Peng Z, Omaruddin R, Bateman E. Stable transfection of *Acanthamoeba castellanii*. Biochim Biophys Acta 2005;1743:93–100

34. Bateman E. Expression plasmids and production of EGFP in stably transfected *Acanthamoeba*. Protein Expr Purif 2010;70:95–100

35. Kapust RB, Waugh DS. *Escherichia coli* maltose-binding protein is uncommonly effective at promoting the solubility of polypeptides to which it is fused. Protein Sci 1999;8:1668–74

36. Wilhelmus KR, Osato MS, Font RL, Robinson NM, Jones DB. Rapid diagnosis of *Acanthamoeba* keratitis using calcofluor white. Arch Ophthalmol 1986;104:1309–12.

37. Shaw JA, Mol PC, Bowers B, Silverman SJ, Valdivieso MH, Durán A, et al. The function of chitin synthases 2 and 3 in the *Saccharomyces cerevisiae* cell cycle. J Cell Biol 1991;114:111–23

38. Frisardi M, Ghosh SK, Field J, Van Dellen K, Rogers R, Robbins P, et al. The most abundant glycoprotein of amebic cyst walls (Jacob) is a lectin with five Cys-rich, chitin-binding domains. Infect Immun 2000;68:4217–24

39. Smith DB. Generating fusions to glutathione S-transferase for protein studies. Methods Enzymol 2000;326:254–70

40. Chen L, Orfeo T, Gilmartin G, Bateman E. Mechanism of cyst specific protein 21 mRNA induction during *Acanthamoeba* differentiation. Biochim Biophys Acta 2004;1691:23–31

41. Chatterjee A, Ghosh SK, Jang K, Bullitt E, Moore L, Robbins PW, Samuelson J. Evidence for a “wattle and daub” model of the cyst wall of entamoeba. PLoS Pathog 2009;5:e1000498

42. Chatterjee A, Carpentieri A, Ratner DM, Bullitt E, Costello CE, Robbins PW, et al. *Giardia* cyst wall protein 1 is a lectin that binds to curled fibrils of the GalNAc homopolymer. PLoS Pathog 2010;6:e1001059

43. Orlean P. Architecture and biosynthesis of the *Saccharomyces cerevisiae* cell wall. Genetics 2012;192:775–818

44. Lampugnani ER, Khan GA, Somssich M, Persson S. Building a plant cell wall at a glance. J Cell Sci 2018; 131

45. Altschul SF, Madden TL, Schäffer AA, Zhang J, Zhang Z, Miller W, et al. Gapped BLAST and PSI-BLAST: a new generation of protein database search programs. Nucleic Acids Res 1997;25:3389–402

46. Aurrecoechea C, Barreto A, Brestelli J, Brunk BP, Caler EV, Fischer S, et al. AmoebaDB and MicrosporidiaDB: functional genomic resources for Amoebozoa and *Microsporidia* species. Nucleic Acids Res 2011;39:D612–9

47. Cabib E, Arroyo J. How carbohydrates sculpt cells: chemical control of morphogenesis in the yeast cell wall. Nat Rev Microbiol 2013;11:648–55

48. Etienne-Manneville S, Hall A. Rho GTPases in cell biology. Nature 2002;420:629–35

49. Sinclair R, Rosquete MR, Drakakaki G. Post-Golgi Trafficking and Transport of Cell Wall Components. Front Plant Sci. 2018;9:1784

50. Oda Y. Emerging roles of cortical microtubule-membrane interactions. J Plant Res 2018;131:5–14.

51. Chebli Y, Geitmann A. Cellular growth in plants requires regulation of cell wall biochemistry. Curr Opin Cell Biol 2017;44:28–35

52. Maeda H, Ishida N. Specificity of binding of hexopyranosyl polysaccharides with fluorescent brightener. J Biochem 1967;62:276–8

53. Meents MJ, Watanabe Y, Samuels AL. The cell biology of secondary cell wall biosynthesis. Ann Bot 2018;121:1107–25

54. Jensen T, Barnes WG, Meyers D. Axenic cultivation of large populations of *Acanthamoeba* castellanii (JBM). J Parasitol 1970;56:904–6

55. Neff RJ, Ray SA, Benton WF, Wilborn M. Induction of synchronous encystment (differentiation) in *Acanthamoeba* sp. Methods Cell Biol 1964;1:55–83

